# Predicting TCR-pMHC Binding by Reinforcement Learning

**DOI:** 10.64898/2026.01.05.697685

**Authors:** Jingxiang Lang, Chungong Yu, Ngoc Hieu Tran, Chao Peng, Qingyang Lei, Haiming Qin, Li Yang, Yi Zhang, Dongbo Bu, Ming Li

## Abstract

The binding between T cell receptors (TCRs) and peptide-major histocompatibility complexes (pMHCs) is fundamental to the immune system’s ability to recognize and eliminate pathogens. Accurate prediction of TCR-pMHC interactions holds significant promise for advancing cancer immunotherapy, vaccine design, and autoimmune disease research. However, existing approaches often treat the sequences, structures, and functions of TCRs, peptides, and MHC molecules in isolation, neglecting their interdependencies and hence limiting the prediction accuracy. In this study, we present ProTCR, a novel approach that integrates sequence, structural, and functional information within a reinforcement learning framework, offering a new paradigm for predicting TCR-pMHC binding. The reinforcement learning optimization enables ProTCR to generate TCR-pMHC sequences with enhanced binding propensity, thereby improving prediction accuracy. On benchmark datasets such as IEDB and VDJdb, ProTCR achieves an AUROC of 0.75, outperforming state-of-the-art methods by 32.7%, while offering interpretable insights into the structural and sequence determinants of binding. We further validate ProTCR using TCRs and neoantigens derived from a cervical cancer patient via proteogenomic profiling. Our analysis reveals a strong correlation between T cell clonal expansion and ProTCR-predicted TCR-peptide binding scores, supporting the biological relevance of the model. Additionally, ProTCR demonstrates robust performance in predicting SARS-CoV-2 TCR-pMHC complexes and generating MHC-specific peptides with potential applications in peptide-based immunotherapies. Collectively, these findings establish ProTCR as a powerful and interpretable tool for TCR-pMHC binding prediction, with broad utility across immunology research and translational applications.

## 1 Introduction

T cell receptors (TCRs) are heterodimeric proteins expressed on the surface of T cells that specifically recognize and bind to peptides presented by major histocompatibility complex (MHC) proteins, thereby mediating T cell activation and initiating adaptive immune responses against foreign antigens [1, 2, 3] (Fig. 1A). TCR-based immunotherapies leverage this natural immune mechanism by engineering TCRs to enhance their affinity and specificity for target antigens, enabling the precise elimination of pathogen-infected or malignant cells. These therapies offer personalized treatment strategies for a variety of diseases, including leukemia and lymphoma [4, 5, 6]. Therefore, accurate prediction of TCR binding to peptide-MHC (pMHC) complexes is essential to elucidate T cell-mediated immune recognition mechanisms and to optimize the design and application of TCR-based immunotherapies [7].

**Fig. 1.**
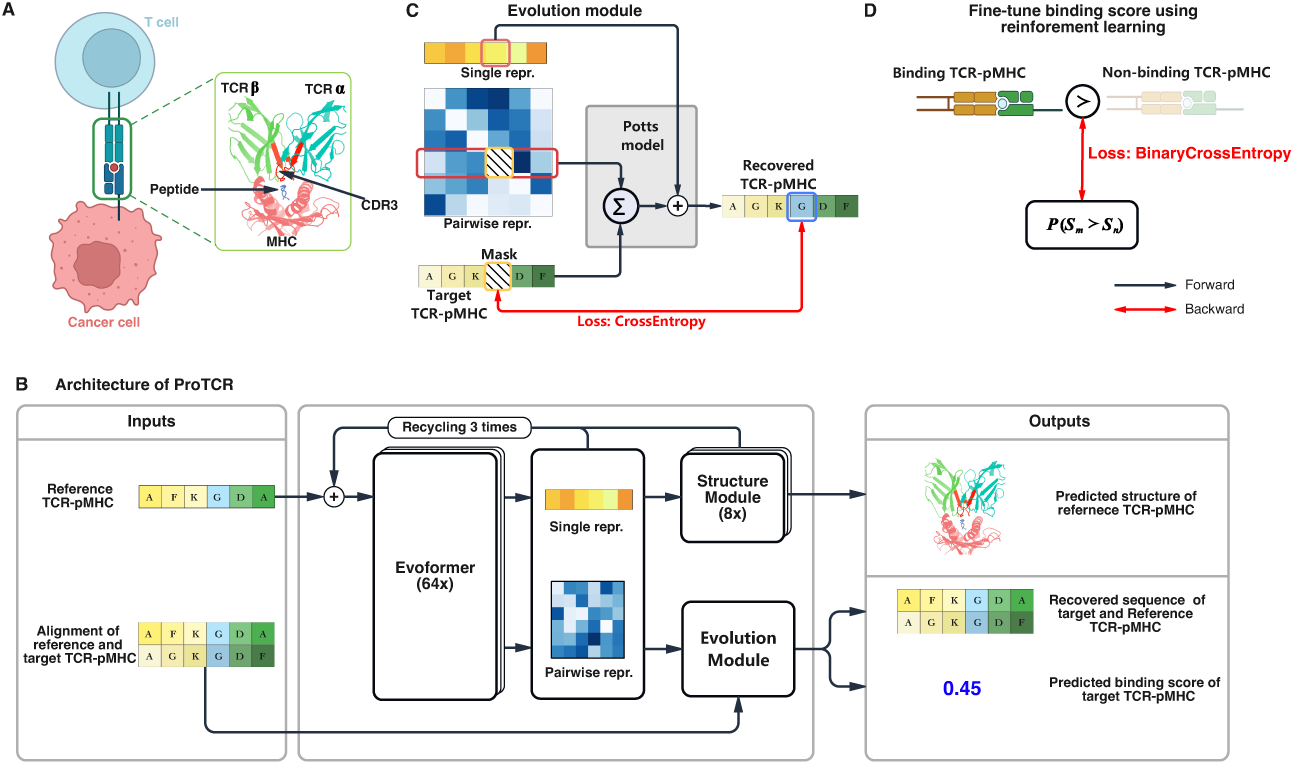
Overview of the ProTCR architecture. **A.** Structural schematic of a TCR-pMHC com-plex, highlighting key interfaces between the TCR, peptide, and MHC molecule. **B.** The architecture of ProTCR approach. ProTCR predicts the probability of a target TCR-pMHC forming a stable interaction through a three-step process: 1) the target TCR-pMHC sequence is searched against a reference database that comprises 103 ground-truth TCR-pMHC complexes with experimentally resolved 3D structure to identify homologous references; 2) the sequence of the identified reference complex is fed into a pre-trained Evoformer to extract single and pairwise representations; and 3) a sequence alignment between the target and reference is fed into the Evolution module to predict the binding score of the target TCR-pMHC complex and to recover the target sequence. In addi-tion, ProTCR also predicts the structure of the target TCR-pMHC complex. The predicted binding score, recovered sequence, and predicted structure are jointly used to calculate the loss function for training ProTCR. **C.** The Evolution module feeds the single and pairwise representations of the ref-erence and target TCR-pMHC complexes into a Potts model. During training, certain residues in the target TCR-pMHC sequence are masked, and the Evolution module attempts to recover them. The cross-entropy between the original and recovered residues is calculated and incorporated into the loss function. **D.** The reinforcement learning strategy used by ProTCR to fine-tune binding scores. Specif-ically, ProTCR employs a Direct Policy Optimization (DPO) strategy to train the scoring model, encouraging it to assign higher scores to binding TCR-pMHC complexes than to non-binding ones. ProTCR measures the ranking of TCR-pMHC complexes derived from the predicted binding scores using binary cross-entropy, which is incorporated as a component of the overall loss function for training.

**Fig. 2.**
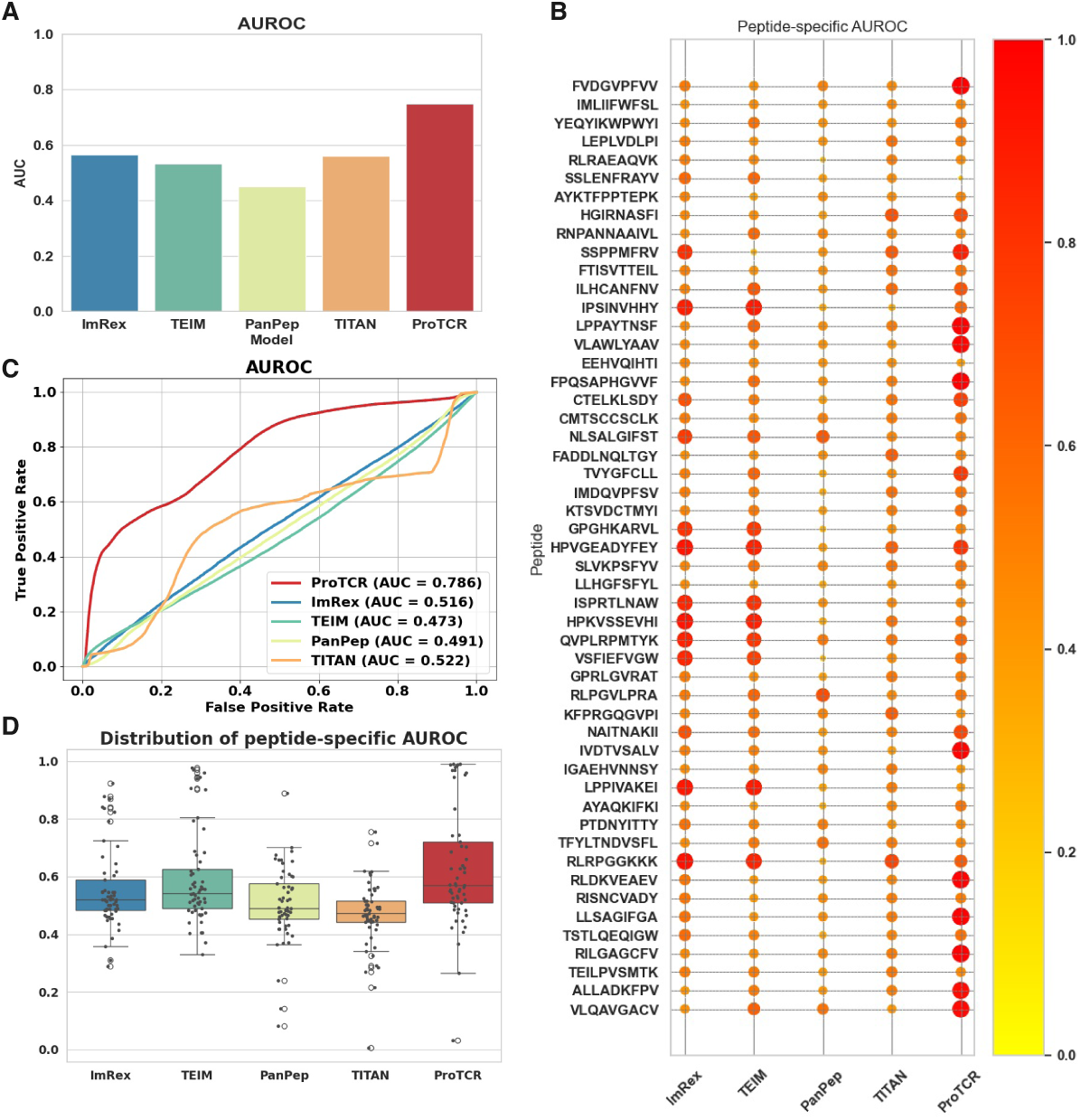
Evaluation of TCR-pMHC binding prediction tools. **A.** AUROC of five prediction models: ImRex, TEIM, PanPep, TITAN, and ProTCR. The average AUROC scores were obtained from five-fold cross-validation on a dataset comprising 173,067 ground-truth TCR-pMHC complexes and 4,651,951 negative samples. ProTCR achieved the highest AUROC score, significantly outper-forming the other models. **B.** Peptide-specific AUROC scores on Subset-1. Darker and larger circles indicate higher AUROC values. ProTCR achieved the highest AUROC for 21 out of 51 peptides, including notable examples such as the SARS-CoV-2 Spike peptide FPQSAPHGVVF (AUC = 0.996), the SARS-CoV-1 epitope VLAWLYAAV (AUC = 0.979), and the Influenza A virus epitope CTELKLSDY (AUC = 0.758). Complete AUROC scores for all peptides are provided in Supplementary Data 1. **C.** ROC curves of the five prediction models on Subset-4. The ROC curves for the remaining four subsets are provided in Supplementary Fig. 1. **D.** Distributions of peptide-specific AUROC scores for Subset-1. The distributions for the remaining four subsets are provided in Supplementary Fig. 2. (ROC: Receiver Operating Characteristic; AUROC: Area Under the Receiver Operating Character-istic Curve).

**Fig. 3.**
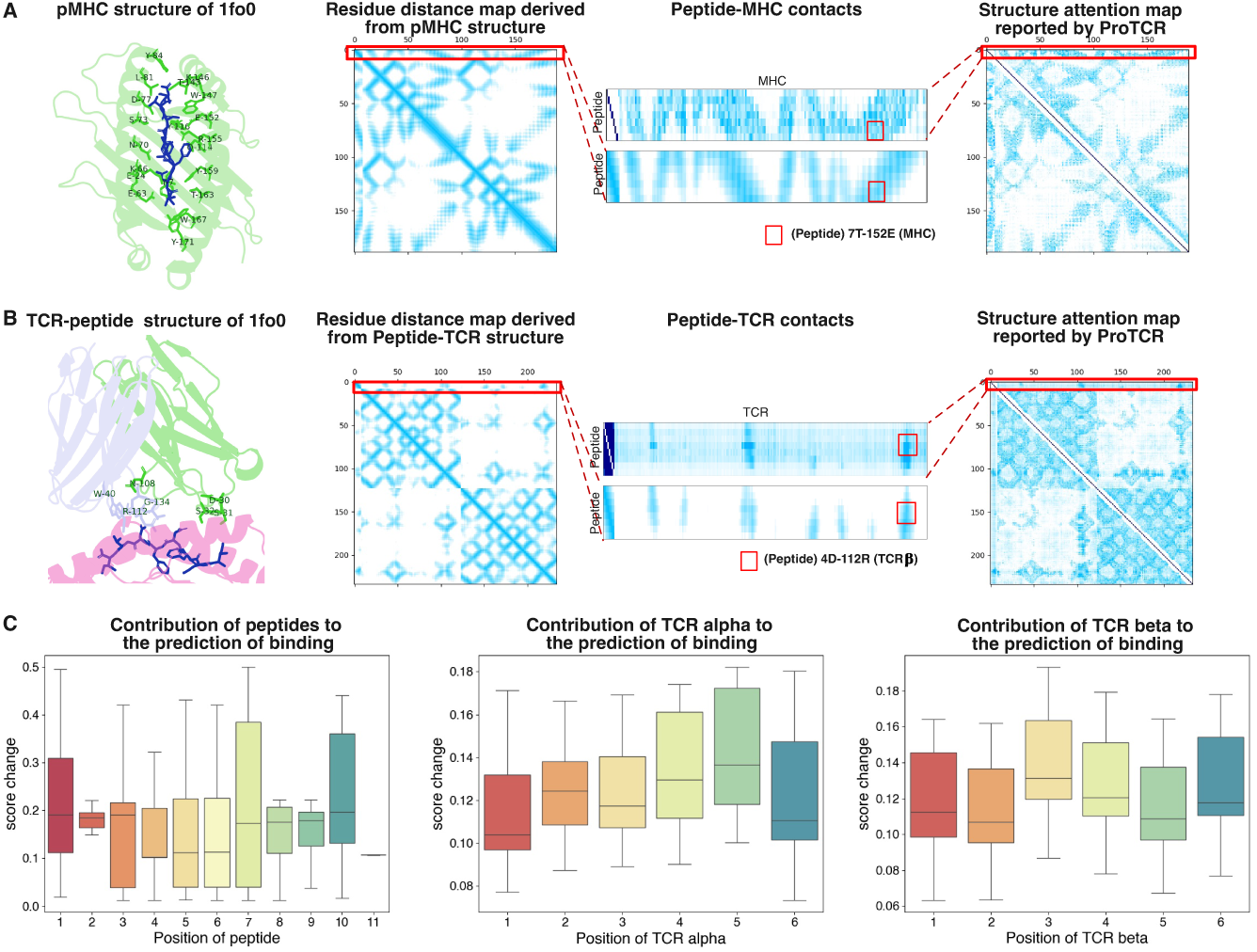
Structural interpretation of TCR-pMHC binding predictions by ProTCR. **A.** Comparison of the inter-residue distance map derived from the pMHC structure of 1fo0 with the structure attention map reported by ProTCR. The peptide and the MHC molecule are shown in blue and green, respectively. The inter-residue distance between the peptide and the MHC molecule perfectly matches the corresponding structure attention reported by ProTCR. **B.** Comparison of the inter-residue distance map derived from the TCR-peptide structure of 1fo0 and the structure attention map reported by ProTCR. The TCR*α* and *β* structures are shown in grey and green, respectively. The inter-residue distance between the peptide and the TCR also perfectly matches the corresponding structure attention reported by ProTCR. **C.** Contributions of residues in the peptide, TCR*α* and *β* chains to predicted binding scores. We assessed the contribution of individual residues by analyzing the effects of point mutations on predicted binding scores. Mutations at peptide positions 1, 2, 3, 7, 8, and 9/10 led to substantial changes, highlighting their critical role in TCR-pMHC binding. In addition, central residues of both TCR*α* and *β* exhibited relatively greater influence on the predictions, implying their importance in TCR-peptide recognition.

A wide range of computational approaches have been developed to predict TCR-pMHC binding by leveraging different sequence, structural, and functional features. Some methods analyze TCR sequence clusters (e.g., TCRdist [8], DeepTCR [9], GIANA [10], iSMART [11], GLIPH [12], TCRGP [13], TCRex [14]); others focus on the characteristics of specific peptides (e.g., MixTCRpred [15], NetTCR-2.0 [16], DeepAIR [17]); and some estimate the binding affinity between TCRs and peptides (e.g., IMRex [18], PanPep [19], TEIM-Res [20], TITAN [21], pMTnet [22], ERGO-II [23]). Confidence metrics from structure predictors like AlphaFold [24, 25] and RoseTTAFold [26], such as pLDDT, pTM, and ipTM, are also used to predict whether TCRs can bind to peptides and MHCs even though they were not trained on this task [27]. Despite these advances, existing approaches still suffer from several limitations, as they often utilize only partial information from the TCR, peptide, and MHC components. For example, many of these methods focus solely on the complementarity-determining region 3 (CDR3) of TCR β chain only [28, 29], whereas TCR-pMHC binding is fundamentally the result of all four chains acting in concert, and neglecting the other three chains greatly limits the prediction accuracy [30, 31, 32]. In addition, most existing approaches primarily focus on sequence features, with limited incorporation of structural properties in TCR-pMHC binding prediction, partly due to the scarcity of high-quality structural data. Furthermore, the limited availability of experimentally validated TCR-pMHC interactions poses challenges such as increased risk of overfitting and reduced generalizability to unseen peptides [33, 34]. Moreover, current methods generally fail to fully utilize the co-evolution information of the TCR-pMHC complex [35, 36], despite its crucial role in accurate binding prediction.

To overcome the limitations of existing methods, we propose ProTCR, a new reinforcement learning approach that integrates sequence, structural, and functional information of TCR-pMHC complexes to model and predict TCR-pMHC binding (Fig. 1). ProTCR consists of three key components. The first is an evolution module, designed to reconstruct homologous sequences of a reference TCR-pMHC complex. This enables ProTCR to learn informative sequence representations, including single representations that capture residue-specific amino acid preferences and pairwise representations that encode co-evolutionary relationships between residue pairs [37, 38]. The second is a structure module that reconstructs the structure of reference TCR-pMHC complexes [39]. This module further optimizes the sequence representations and provides structural insights to assess the likelihood of TCR, peptide, and MHC binding, as well as to identify potential binding sites [40]. The third component is a binding score function that employs a learn-to-rank technique to distinguish binding TCR-pMHC complexes from constructed negative (non-binding) samples. By leveraging reinforcement learning [41, 42], ProTCR is guided to generate sequences with higher binding propensity, thereby improving the accuracy of binding prediction. Unlike existing methods that focus primarily on CDR3 regions, ProTCR utilizes full-length TCR sequences, fully exploiting the complete information encoded in TCRs to enhance both predictive performance and biological interpretability.

We evaluated ProTCR on the IEDB and VDJdb datasets and found that it achieved an AUROC of 0.75 on unseen peptides, substantially outperforming existing state-of-the-art methods. In addition, ProTCR identified potential binding sites and sequence motifs, offering interpretability and facilitating an in-depth understanding of the molecular determinants of TCR-pMHC interactions. We further assessed ProTCR’s capability to generate MHC-specific binding peptides, demonstrating its potential utility in the design of peptide-based immunotherapies. Finally, we applied ProTCR to the ImmuneCODE dataset to investigate viral immune recognition, and to a proteogenomic dataset from a cervical cancer patient to evaluate neoantigen-specific TCR responses. In the latter case, we observed a strong correlation between predicted TCR-peptide binding scores and T cell clonal expansion, supporting the biological relevance of ProTCR’s predictions. Collectively, these findings establish ProTCR as a powerful and interpretable framework for TCR-pMHC binding prediction and broader applications in immunological research.

## 2 Results

### 2.1 Overview of the ProTCR architecture

ProTCR takes as input the full-length sequences of the target TCR-pMHC components, including the TCR α and β chains, the peptide, and the MHC molecule, and predicts the probability that these components form a stable complex. To achieve this, ProTCR compares the target TCR-pMHC against a curated set of 103 ground-truth

TCR-pMHC complexes with experimentally resolved 3D structures, referred to hereafter as *reference TCR-pMHCs*. Fig. 1B shows the three main steps of ProTCR. First, ProTCR identifies homologous references by searching the target sequence against the reference database. Second, the identified reference TCR-pMHC sequence is fed into a pre-trained Evoformer to extract single and pairwise representations. Third, these representations, along with a sequence alignment between the target and reference, are passed into the evolution module to generate a binding score that estimates the likelihood of the target TCR-pMHC forming a stable complex. In addition, ProTCR also predicts the structure of the target TCR-pMHC complex.

The strength of ProTCR lies in its training process, which jointly trains its key components, Evoformer, the structure module, and the evolution module, to capture information about sequence, structure, and evolutionary constraints (Supplementary Figure 1). Specifically, Evoformer is trained to extract both single and pairwise representations of reference TCR-pMHCs efficiently. The structure module is designed to reconstruct the structures of reference TCR-pMHC complexes, further refining the single and pairwise representations while providing insights into TCR-pMHC binding. The evolution module is trained to recover the homologous sequences of reference TCR-pMHCs, enabling ProTCR to explore the evolutionary constraints associated with these homologues. Based on the refined representations, the evolution module also predicts a binding score for the target TCR-pMHC. To achieve this goal, the evolution module is trained using the Direct Preference Optimization (DPO) strategy [43], which encourages the model to assign higher scores to ground-truth TCR-pMHC complexes than to non-binding ones (Fig. 1D). This reinforcement learning approach significantly enhances ProTCR’s accuracy in binding prediction.

### 2.2 Evaluation of ProTCR for TCR-pMHC binding prediction

We evaluated ProTCR by assessing its ability to distinguish ground-truth TCR-pMHC complexes from non-binding (negative) pairs. A total of 196,452 experimentally validated TCR-pMHC complexes were compiled from the IEDB [44] and VDJdb [45] databases. To ensure a fair comparison, negative samples were generated using the *random epitope* approach [46], consistent with the method employed by TEIM [20]. Specifically, TCRs were extracted from the ground-truth TCR-pMHC complexes and paired with peptides randomly sampled from the complete epitope repertoire. Due to the random pairing, these artificial TCR-pMHC combinations have a low probability of binding and are therefore considered negative samples [20]. The ratio of ground-truth to negative samples was controlled at 1: 5. Given this class imbalance, the prediction performance was evaluated using the area under the receiver operating characteristic curve (AUROC). Additionally, TCR-pMHCs that could not be perfectly aligned with any reference complex were excluded, resulting in a final dataset comprising 173,067 ground-truth complexes and 4,651,951 negative samples.

To prevent the overlap between training and test sets, the peptides were divided into five non-overlapping groups and the corresponding TCR-pMHCs were partitioned into five subsets accordingly (hereafter referred to as Subset-1∼5). A five-fold cross validation was performed, with each subset used as the test set while the remaining four subsets served as the training data. The average AUROC score across the five validation rounds was used as the final evaluation metric (See Methods for details). This strategy ensures that no peptides are shared between training and test sets, avoiding information leakage. More importantly, this experimental setup also simulates the prediction of TCR-pMHC interactions involving previously unseen neoantigens, thus offering a more objective evaluation across diverse peptide repertoires.

Fig. B3 shows the evaluation performance of ProTCR in comparison with four existing methods, including ImRex [18], TEIM [20], PanPep [19], and TITAN [21].

ProTCR achieved an average AUROC of 0.75, significantly outperforming the other methods (Fig. B3A; ImRex: 0.57, TEIM: 0.53, PanPep: 0.45, TITAN: 0.56). The corresponding ROC curves are shown in Fig. B3C and Supplementary Figure 1. These results indicate that ProTCR is at least 32% more accurate in distinguishing ground-truth TCR-pMHC complexes from non-binding pairs.

In addition, we also compared ProTCR with AlphaFold3, which has demonstrated strong performance in predicting TCR-pMHC binding despite not being explicitly trained for this task [27]. Due to computational constraints, we randomly selected 1,000 entries from the 10x single-cell sequencing library, each containing complete TCR α, TCR β, peptide, and MHC sequences, as our test set. To avoid data leakage, ProTCR predictions for each test sample were generated using models that excluded that sample from training. ProTCR outperformed AlphaFold3 in top-k precision (Supplementary Figure 3), and notably, there was minimal overlap between the top-k predictions of the two models.

To investigate the impact of peptides on TCR-pMHC binding, we examined the peptide-specific prediction accuracy. Specifically, each test subset was further divided into groups in which all TCR-pMHC complexes shared the same peptide, and the AUROC was then calculated for each group (hereafter referred to as peptide-specific AUROC). Groups containing fewer than 10 TCR-pMHC complexes were excluded to ensure the statistical reliability of the evaluation. For Subset-1, the data was divided into 66 peptide-specific groups. As shown in Figure B3D, ProTCR achieved an average peptide-specific AUROC of 0.53, outperforming the other methods. Similar results were observed across the remaining four subsets (Supplementary Figure 2). Notably, ProTCR demonstrated the highest predictive accuracy for 21 out of 51 peptides (Fig. B3B, see Supplementary Data 1 for more details), including the SARS-CoV-2 Spike peptide FPQSAPHGVVF (AUC = 0.996), the SARS-CoV-1 epitope VLAWLYAAV (AUC = 0.979), and the Influenza A virus epitope CTELKLSDY (AUC = 0.758). Overall, these results collectively validate ProTCR’s capacity to generalize across phylogenetically diverse pathogenic targets while maintaining high predictive fidelity.

### 2.3 Structural insights into TCR-pMHC binding uncovered by ProTCR’s attention mechanism

In addition to predicting a binding score, ProTCR provides structural insights to support the interpretation of its predictions. Previous studies have elucidated the structural mechanisms underlying protein complex formation, highlighting the importance of compatible structural conformations between interacting partners and the presence of key interface residue pairs that mediate binding [47]. The structure module of ProTCR employs a self-attention mechanism to capture the local structural context surrounding each residue. By analyzing the resulting attention matrices, we can identify key interface residue pairs and characterize their distribution based on residue type, position, and interaction class. Such an analysis provides meaningful structural insights into the formation of TCR-pMHC complexes.

For instance, Fig. B4A shows the inter-residue distance map derived from the pMHC structure of 1fo0 together with the structure attention map reported by ProTCR. The attention map perfectly resembled the ground-truth distance map across nearly all residue pairs, including the residues within the MHC and peptide, as well as the interface residues between them, e.g., the contact between peptide residue 7T and MHC residue 152E. Similar patterns were observed for residue contacts between the TCR and the peptide (Fig. B4B). These findings confirm ProTCR’s ability to recover meaningful structural interactions.

To assess the contributions of residues in the peptide and TCRα and β chains to TCR-pMHC binding, we employed ProTCR to perform *in silico* mutagenesis on both the TCR and peptide sequences. Specifically, we first simulated residue perturbation using a zero-vector scanning strategy, where the embedding of a target residue was replaced with a zero vector [22], and then measured the change in the predicted binding score. We analyzed CDR3 regions and peptides across 104 TCR-pMHC complexes and found that mutations at peptide residues 1, 2, 3, 7, 8, and 9/10 resulted in significant decreases in the predicted binding scores (Fig. B4C). Notably, this observation aligns with the previously identified key interface residue pair (peptide 7T-152E MHC, Fig. B4A). We further performed segment-wise perturbation analysis on CDR3α and CDR3β by dividing each chain into six equal-length segments and evaluating the impact of mutations within each segment on the predicted binding scores. As shown in Fig. B4C, mutations in central regions of CDR3β (segments 3 and 4) and CDR3α (segments 4 and 5) led to the most substantial decreases in binding scores, highlighting the critical role of these central segments in TCR-pMHC recognition. These results demonstrate that ProTCR accurately captures residue-level contacts and identifies key interface residues, supporting the interpretation of its TCR-pMHC binding predictions.

### 2.4 Generating MHC-specific peptides for applications in peptide-based immunotherapy

MHCs recognize and bind to specific peptides with distinct residue preferences. These preferences, often described as sequence motifs, define the specificity of MHC-peptide interactions and play a crucial role in antigen presentation by the immune system [48]. Here we assessed whether ProTCR could generate MHC-specific peptides. To do this, we masked the peptides from ground-truth TCR-pMHC complexes and attempted to recover them using ProTCR, based solely on the corresponding TCR and MHC sequences. The recovered peptides are then compared to the original ground-truth peptides to assess reconstruction accuracy.

Fig. 4 shows the sequence motifs of peptides recognized by three representative MHC alleles, including A0201, B0702, and C0701, acquired from 8372, 6747, and 891 ground-truth TCR-pMHC complexes, respectively. These peptides exhibit substantial residue preferences: MHC A0201 favors peptides with conserved residues 2LEU and 9LEU/VAL; B0702 prefers 2PRO and 9LEU; and C0701 tends to recognize 1ARG, 2ARG, and 9LEU/TYR/PHE. Remarkably, the peptides recovered by ProTCR, based on the corresponding MHC and TCR, exhibit sequence motifs that closely mirror those of the ground-truth peptides, including nearly identical conserved residues. We repeated this analysis across 12 additional MHC alleles and observed similar results (Supplementary Fig. 4). These findings indicate that ProTCR can effectively learn sequence patterns and, more importantly, generate biologically meaningful peptides that are specific to individual MHC alleles. This not only provides insights into the determinants of TCR-pMHC binding specificity but also highlights the model’s potential for applications in the design of MHC-specific peptides for immunotherapy and vaccine development.

**Fig. 4.**
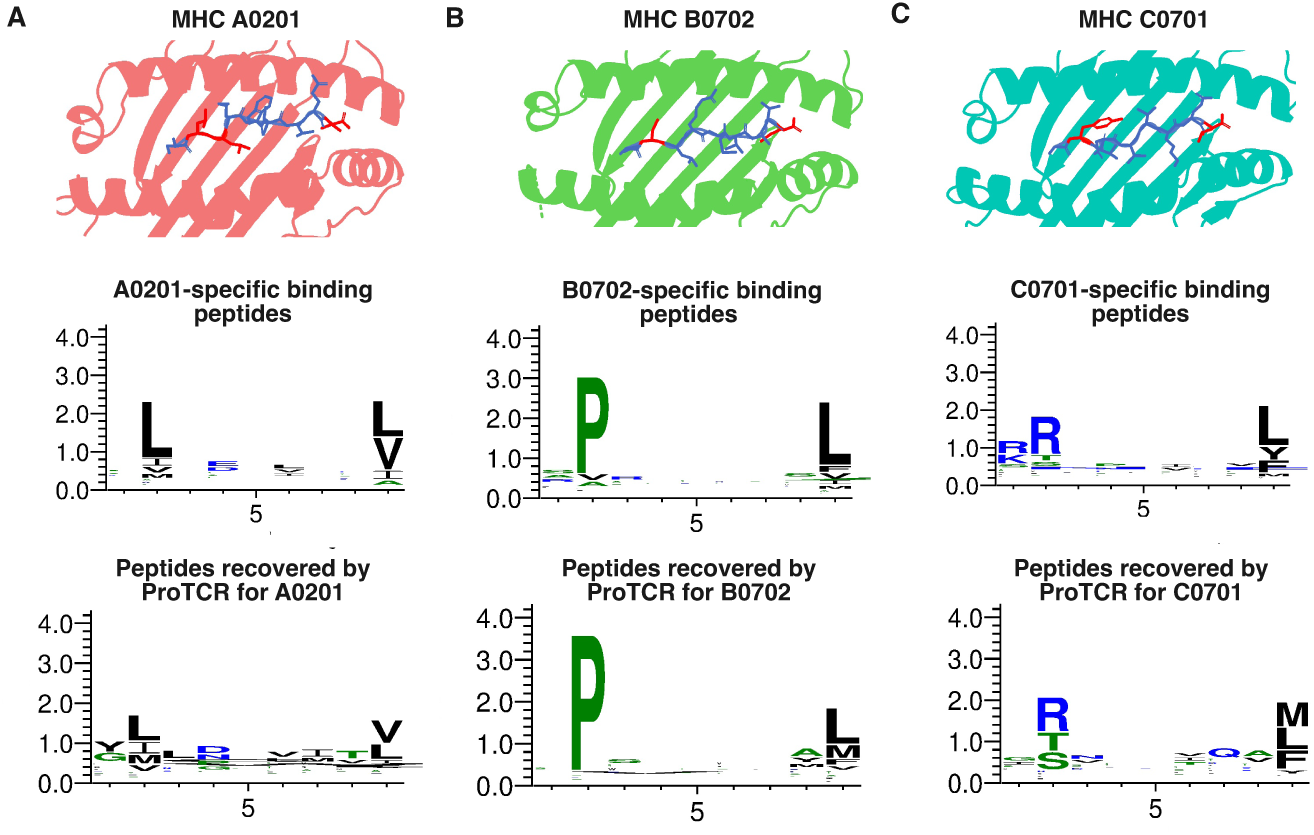
Sequence motifs of MHC-bound peptides (ground-truth) and peptides generated by ProTCR for three representative MHC alleles: A0201, B0702**, and** C0701**. A.** Sequence motif of peptides recognized by MHC A0201 with conserved residues 2LEU and 9LEU/VAL. **B.** Sequence motif of peptides recognized by MHC B0702 with conserved residues 2PRO and 9LEU. **C.** Sequence motif of peptides recognized by MHC C0702 with conserved residues 1ARG, 2ARG, and 9LEU/TYR/PHE. In the first row, the peptides are shown in blue, with conserved residues highlighted in red. The middle row displays the sequence motifs of MHC-bound peptides extracted from the ground-truth TCR-pMHC complexes, while the bottom row shows the motifs of peptides recovered by ProTCR for the corresponding MHC alleles.

### 2.5 Application of ProTCR for predicting SARS-CoV-2 TCR-pMHC complexes

In this section, we demonstrate the application of ProTCR for predicting SARS-CoV-2 TCR-pMHC complexes using the ImmuneCODE [49] dataset (Fig. 5A). This dataset profiles TCR repertoires from over 1,400 individuals exposed to or diagnosed with COVID-19 and includes millions of TCR sequences with high-confidence virus-specific annotations. After rigorous preprocessing and deduplication, we curated a large-scale dataset of 1,410,560 TCR-peptide pairs spanning 246 distinct viral peptides (see Meth-ods), which serve as the ground-truth samples in our study. In addition, we generated negative samples by randomly shuffling the ground-truth pairs at a 1:5 ratio of pos-itive to negative samples. We then assessed whether the pre-trained ProTCR model could effectively distinguish true ImmuneCODE TCR-peptide pairs from these artifi-cial negatives. To ensure a fair evaluation, all peptides present in ProTCR’s training set were excluded from this analysis.

**Fig. 5.**
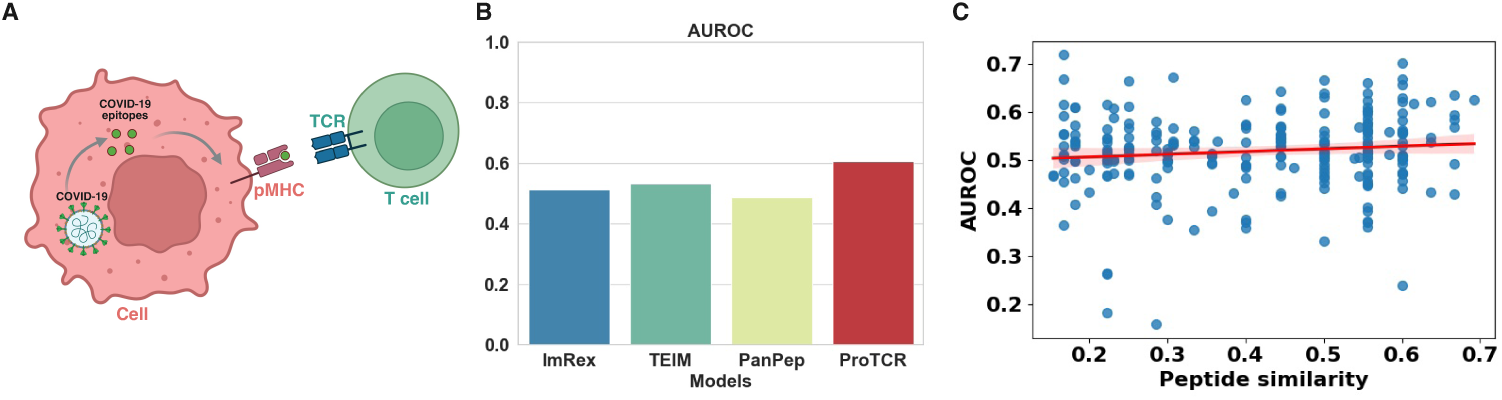
Evaluation of ProTCR for predicting SARS-CoV-2 TCR-pMHC complexes. **A.** Schematic illustration of antigen presentation and T cell recognition in a SARS-CoV-2-infected cell. **B.** AUROC of four prediction models: ImRex, TEIM, PanPep, and ProTCR. ProTCR achieved the highest AUROC score of 0.606, while the other models yielded AUROCs close to 0.50. **C.** Correlation between AUROC scores and the similarity of SARS-CoV-2 peptides to those in the ProTCR training set. The low Pearson correlation coefficient of 0.11 indicates that ProTCR reliably predicts TCR-peptide binding even for peptides highly dissimilar to those seen during training.

As shown in Fig. 5B, ProTCR achieved an AUROC of 0.606 on this challenging dataset, whereas existing approaches yielded AUROCs close to 0.50. These results highlight ProTCR’s significant improvement in peptide-specific TCR recognition com-pared to existing tools. We further examined whether ProTCR’s performance was influenced by the similarity between SARS-CoV-2 peptides and those seen during training. As shown in Fig. 5C, the Pearson correlation coefficient between AUROC values and peptide similarity is only 0.11, suggesting that ProTCR can reliably pre-dict TCR-peptide binding even when the peptides are highly dissimilar from those in the training set (see Supplementary Data 2 for more details).

### 2.6 ProTCR accurately predicts neoantigen-specific TCRs associated with T cell clonal expansion

TCRs initiate effective immune responses by recognizing pMHC complexes and triggering the clonal expansion of antigen-specific T lymphocytes. It is reasonable to hypothesize that TCRs with higher binding affinity for a given antigen may drive greater clonal expansion of the corresponding T cells [50]. In this section, we investigate whether ProTCR can qualitatively capture the relationship between predicted TCR-pMHC binding affinity and clonal behavior in T cells.

We first performed proteogenomic profiling of a cervical cancer patient and identified a panel of experimentally validated immunogenic neoantigens (Fig. 6A; see Ref. [51] for further details). To ensure robust analysis, we excluded outlier TCRs with fewer than 50 clones. As a result, we obtained a total of 6,338 TCRα sequences, 12,047 TCRβ sequences, and 2 neoantigens (QLYEGKLYY and QLNKIQIKH) that exhibited significantly elevated ELISPOT responses. We then enumerated all possible TCR-peptide pairs and predicted their binding scores using ProTCR. As shown in Fig. 6B, the predicted TCR-peptide binding scores positively correlate with the number of TCR clones. This observation supports the hypothesis that high-affinity TCRs undergo preferential clonal expansion. Notably, the immunodominant peptide QLNKIQIKH exhibited a stronger correlation between ProTCR scores and clonal frequency than QLYEGKLYY, consistent with expectations.

**Fig. 6.**
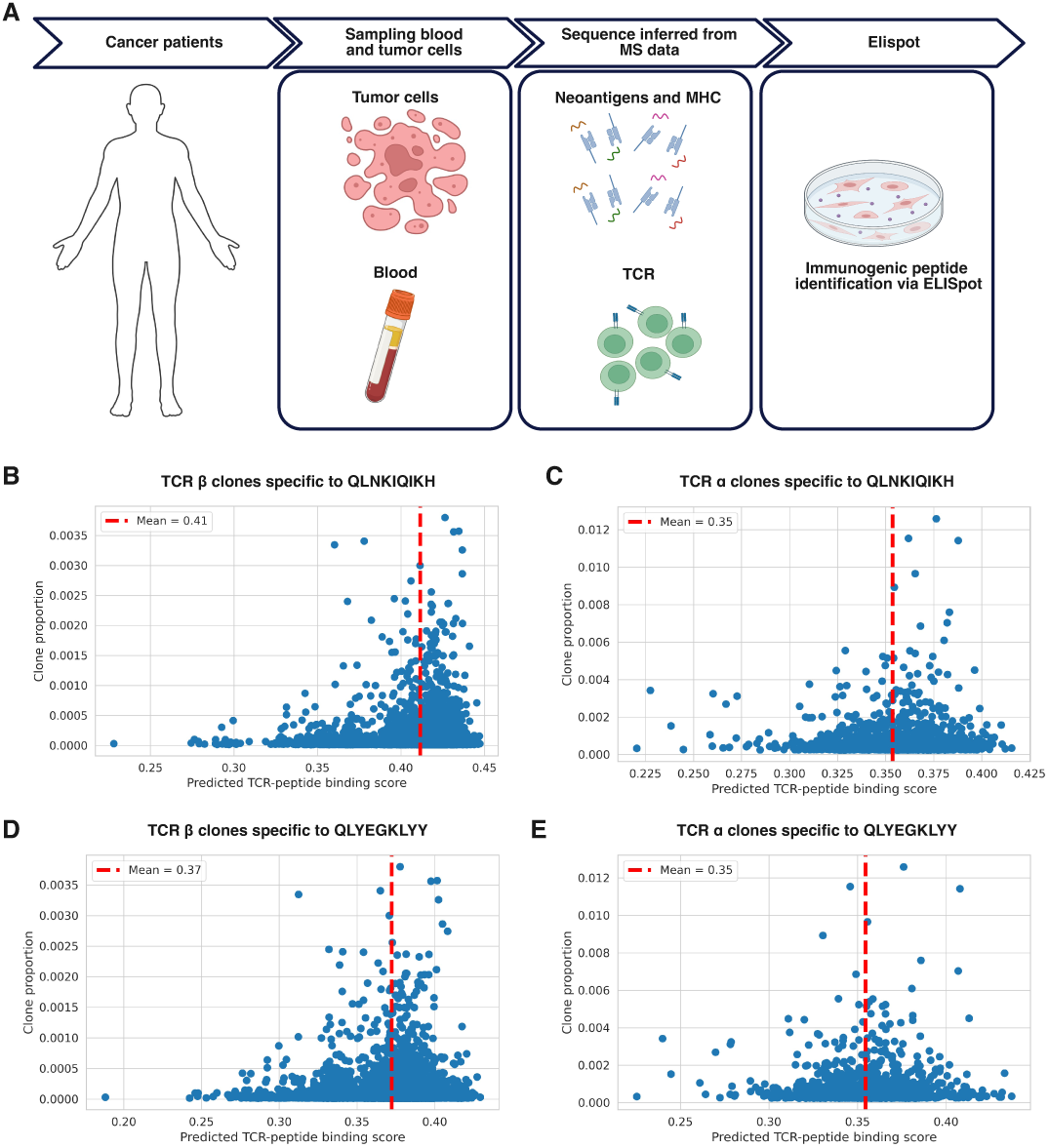
Correlation between ProTCR-predicted TCR-neoantigen binding scores and T cell clonal expansion. **A.** Experimental workflow for acquiring TCRs and neoantigens through proteogenomic profiling of a cervical cancer patient. Tumor tissue and peripheral blood were collected from a patient and subjected to mass spectrometry immunopeptidomics and TCR sequencing to identify neoantigens and TCR repertoires. Candidate neoepitopes were subsequently validated for immunogenicity using the ELISPOT assay. **B-C.** Correlation between the abundance of TCR*β* and TCR*α* clones specific to the peptide QLNKIQIKH and their predicted binding scores by ProTCR. **D-E.** Correlation between the abundance of TCR*β* and TCR*α* clones specific to the peptide QLYEGKLYY and their predicted binding scores by ProTCR.

## 3 Discussion

In this study, we present ProTCR, a unified framework for predicting TCR-pMHC binding by integrating full-length sequence representations, structural embeddings, and reinforcement learning. Unlike existing models that rely solely on handcrafted features of CDR3β sequences, ProTCR incorporates structure prediction and learn-to-rank techniques to deliver state-of-the-art performance, while offering enhanced interpretability and computational efficiency.

ProTCR demonstrates strong generalization to unseen epitopes, including both viral epitopes and cancer neoantigens. Across benchmark datasets such as IEDB, VDJdb, and the ImmuneCODE dataset from COVID-19 patients, ProTCR consistently outperforms current state-of-the-art methods by a large margin. Importantly, its performance remains robust even for peptides with low similarity to the training data, highlighting its potential for real-world applications in immunotherapy. In addition, ProTCR offers structural and residue-level insights into TCR-pMHC interactions through its attention mechanisms and contact-aware representations. Perturbation analyses based on *in silico* mutations reveal key residues within both the peptide and TCR CDR3 regions. Moreover, the model successfully recovers MHC binding motifs and its predictions show strong correlation with T cell clonal expansion, further supporting its biological relevance. Compared to structure-based models that rely on predicted structures, ProTCR offers greater scalability by leveraging precomputed reference embeddings and supporting batched MSA-format inputs. Its flexible architecture enables inference even with partial inputs (e.g., TCR and peptide), making it applicable to a broader range of datasets.

Despite these advancements, some challenges and opportunities remain open for future studies. Unlocking the full clinical potential of TCR-pMHC prediction will require continued efforts to expand datasets, particularly to include a wider diversity of MHC alleles, TCR repertoires, and pathogen-derived epitopes. Further refinement in integrating sequence and structural information, alongside improved model interpretability, will be critical for both clinical translation and mechanistic insight. Future directions may also focus on the incorporation of advanced reinforcement learning strategies and enhanced structural alignment methods, which may improve the detection of rare, cross-reactive, or low-affinity TCR-pMHC interactions.

Overall, the findings presented here establish ProTCR as an effective approach and a powerful tool for predicting TCR-pMHC binding. We anticipate that ProTCR will serve as a valuable tool in advancing cancer immunotherapy, vaccine development, and research into autoimmune diseases.

## 4 Data sets and Methods

### 4.1 Data sets

#### TCR-pMHC data processing

We first collected comprehensive TCR-pMHC binding records from the IEDB [44] and VDJdb [45] databases. The collected data were then filtered and processed according to the following criteria: only records associated with MHC I alleles were retained, and entries lacking peptide information were removed. To construct complete TCR sequences, we utilized the Stitchr tool to assemble CDR3 sequences with their corresponding VJ genes, while filtering out sequences where both the TCR α chain or TCR β chain failed to be reconstructed. After data preprocessing, we obtained 1,372 unique peptides and 196452 valid peptide-TCR binding pairs.

#### pMHC data processing

We first collected pMHC data from NetMHCpan [48] as the training dataset, retaining only records associated with MHC-I alleles. After filtering, we obtained 888034 unique peptides and 897956 valid pMHC binding pairs.

#### Sequence alignment approach

To align binding data with structural data, we collected 104 three-dimensional structures of TCR-pMHC complexes from the RCSB PDB database. The alignment of TCR-pMHC binding data and pMHC binding data to structural data was performed based on the following criteria: TCR α or TCR β Chain Alignment: TCR sequences from PDB structural data were aligned to TCR sequences in the TCR-pMHC dataset using jackhmmer [52].

MHC Sequence Alignment: MHC sequences from PDB structural data were first aligned to MHC sequences in the TCR-pMHC and pMHC datasets using jackhmmer. Alignment results were further filtered to retain only matches with a sequence identity greater than 0.85, ensuring structural consistency.

Peptide Sequence Alignment: Peptide sequences from PDB structural data were aligned to sequences in the TCR-pMHC and pMHC datasets, ensuring that only peptides of identical length were considered.

The TCR-pMHC complex is successfully aligned to the reference TCR-pMHC com-plex if and only if each chain in the target TCR-pMHC complex can be successfully aligned to the reference TCR-pMHC complex.

#### Construction of the five-fold cross-validation dataset

To strictly avoid data leakage in model evaluation, a Five-Fold Cross-Validation Dataset was constructed based on unseen data. The specific steps are as follows: Peptide Clustering: Clustering was performed on 1,372 unique peptides in the cleaned dataset. First, the Levenshtein distance between all antigen sequences was calculated to generate a distance matrix, which served to quantify sequence similarity:

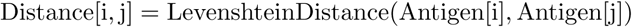

The entry Distance[i,j] in this symmetric matrix holds the calculated Levenshtein distance between antigen i and antigen j, indicating the sequence similarity between them.

K-means clustering [53] was then used on the antigen sequences with the number of clusters set to 500 to maximize inter-cluster differences and balance intra-cluster sample sizes:

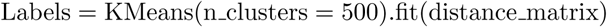

Data Partitioning: Based on clustering results, the peptides were divided into five groups. For each cross-validation fold, four groups were used as the training set, and the remaining group served as the test set, ensuring independence between peptide sequences.

### Generation of negative samples

The direct use of the TCR data collected by external libraries can lead to overesti-mation of results. To avoid this issue, negative samples were generated by randomly pairing TCR and pMHC sequences from the training set. This pairing process ensured that the randomly paired peptides and the original peptide of the TCR did not belong to the same cluster, avoiding potential associations.

### 4.2 Methods

In this study, we developed a model for TCR-pMHC affinity ranking, utilizing multiple sequence alignment (MSA) to generate amino acid embedding vectors and calculating the affinity between different TCR-pMHC sequences based on these embedding vectors. The specific steps are as follows:

#### One-hot encoding of amino acids

For each target TCR-pMHC sequence S*_m_* aligned to the reference in the MSA, the amino acid S*_m,i_* at position i is first converted into one-hot encoding. This process is represented as follows:

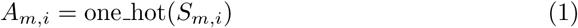

Here, A*_m,i_* denotes the 20−dimensional one-hot vector corresponding to the amino acid at position i in sequence S*_m_*.

#### Calculation of embedding representations

We use the Potts model to compute the embedding representation of the TCR-pMHC sequences, considering the distribution of amino acids at different positions as well as the influence of neighboring amino acids. The calculation formula for the embedding representation h*_m,i_*is given by:

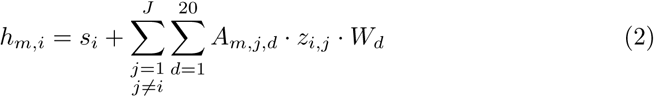

Here, s*_i_* and z*_i,j_* are derived from the single and pairwise representations from Evo-former, while W*_d_* is the weight matrix. The formula effectively integrate the contextual information of the i-th position in the TCR-pMHC sequence with that of other positions.

#### Calculation of Hamiltonian difference between sequences

To compare the affinity differences between the m-th TCR-pMHC sequence and the n-th TCR-pMHC sequence, we calculate their hamiltonian difference δh*_m,n_*:

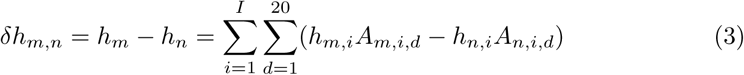

Here, I is the total length of the TCR-pMHC sequences. This method allows us to quantify the differences between two TCR-pMHC sequences in the embedding space.

#### Calculation of ranking of binding affinity

Based on the Hamiltonian difference, we further calculate the affinity ranking probability between the m-th TCR-pMHC sequence and the n-th TCR-pMHC sequence in the given MSA.

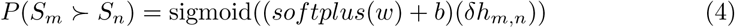

Here, w denotes a linear transformation that maps the Hamiltonian difference to the probability space, b is a constant value of 0.01. The output probability value ranges from 0 to 1, indicating the affinity of the m-th TCR-pMHC sequence compared to the n-th TCR-pMHC sequence.

#### Calculation of amino acid class probability

Finally, we compute the probability that the m-th TCR-pMHC sequence at position i belongs to class amino acid:

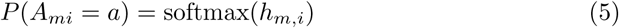

This formula transforms the embedding vector h*_m,i_* into a probability distribution over amino acid categories at the specified position, where a denotes the amino acid type at that position.

Through the above methods, we can accurately calculate the affinity between dif-ferent TCR-pMHC sequences within the context of TCR-pMHC affinity ranking, and further analyze the class probabilities of amino acids. This method provides an effective tool for the structural and functional analysis of TCR-pMHC.

## 5 Data availability

The datasets used for model training and evaluation were compiled from mul-tiple public resources. TCR–pMHC interaction data were integrated from IEDB (http://www.iedb.org/) and VDJdb (https://vdjdb.cdr3.net/), as well as the Pan-Pep benchmark suite (https://github.com/bm2-lab/PanPep). A large-scale, pub-licly available COVID-19 TCR repertoire dataset was obtained from Adap-tive Biotechnologies (https://clients.adaptivebiotech.com/pub/covid-2020). Refer-ence structures of experimentally resolved TCR–pMHC complexes were retrieved from the Protein Data Bank (PDB; https://www.rcsb.org). The comparison with AlphaFold3 was performed using data from the 10X Genomics Datasets (https://www.10xgenomics.com/resources/datasets)

## 6 Code availability

PorTCR is available on Github (https://github.com/bigict/ProTCR)

## Acknowledgements

We thank Shuai Cheng Li for his guidance and assistance with the immunological datasets. We are also grateful to our laboratory colleagues for their consistent encouragement and technical support throughout this study. This work was supported in part by the National Key Research and Development Program of China (2024YFA1306401) [pdflatex,sn-mathphys-num]sn-jnl graphicxmultirowamsmath,amssymb,amsfontsamsthmmathrsfs[title]appendixxcolortextcompmanyfootbo Theorem[theorem]Proposition ExampleRemark Definition [Supplementary Information] Predicting TCR-pMHC Binding by Reinforcement Learning

## Supplementary Information

## 1 Supplementary Methods

**Algorithm 1** ProTCR - Prediction procedure

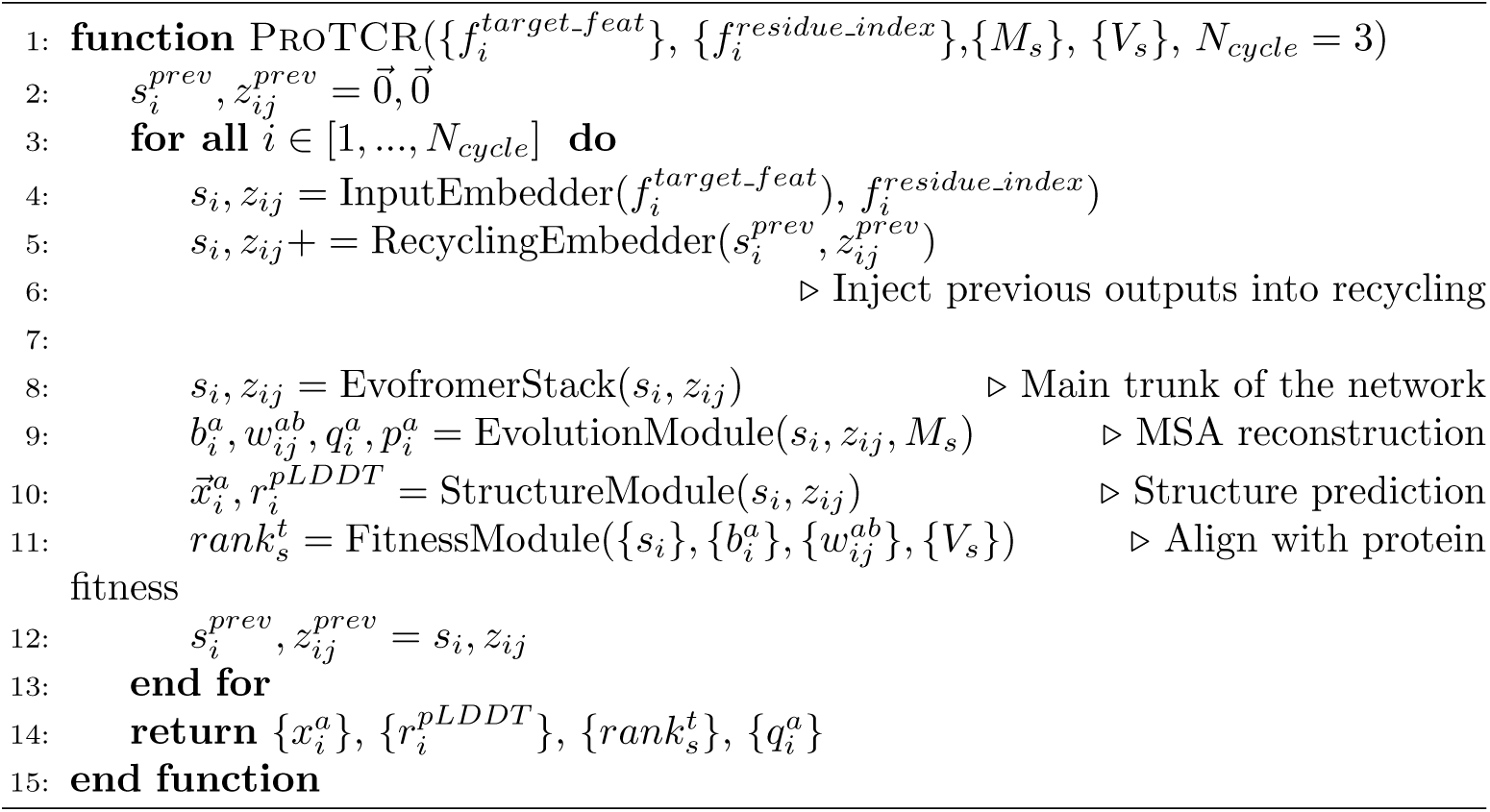

**Algorithm 2** Evolution Module

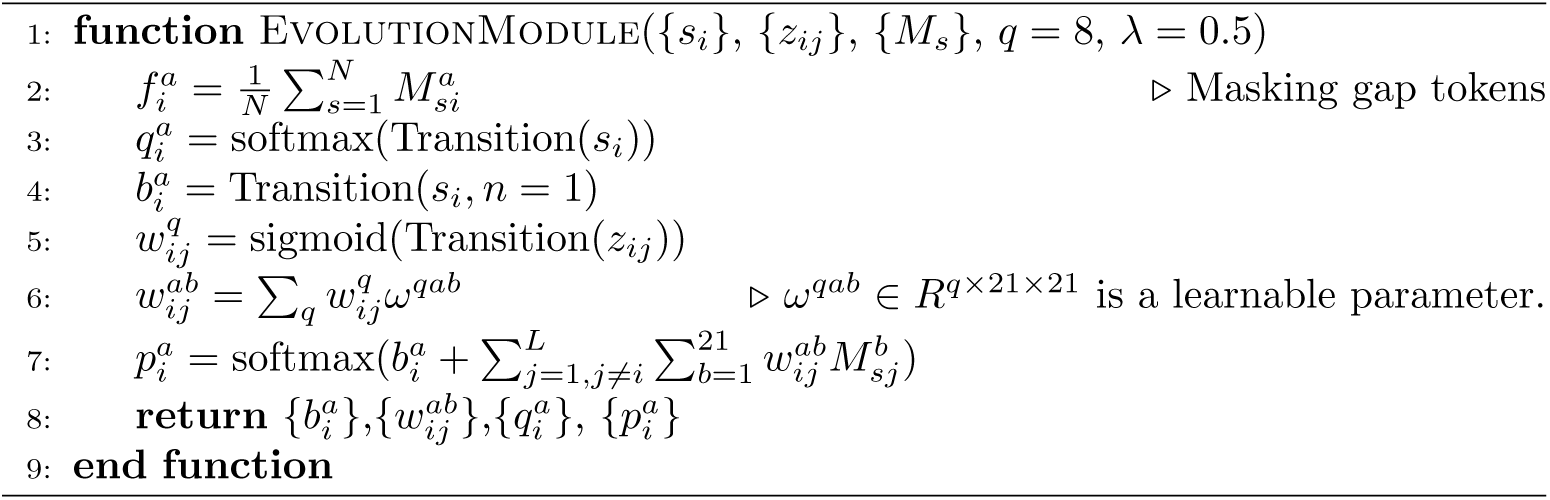

**Algorithm 3** Fitness Module - Align ProTCR with protein fitness

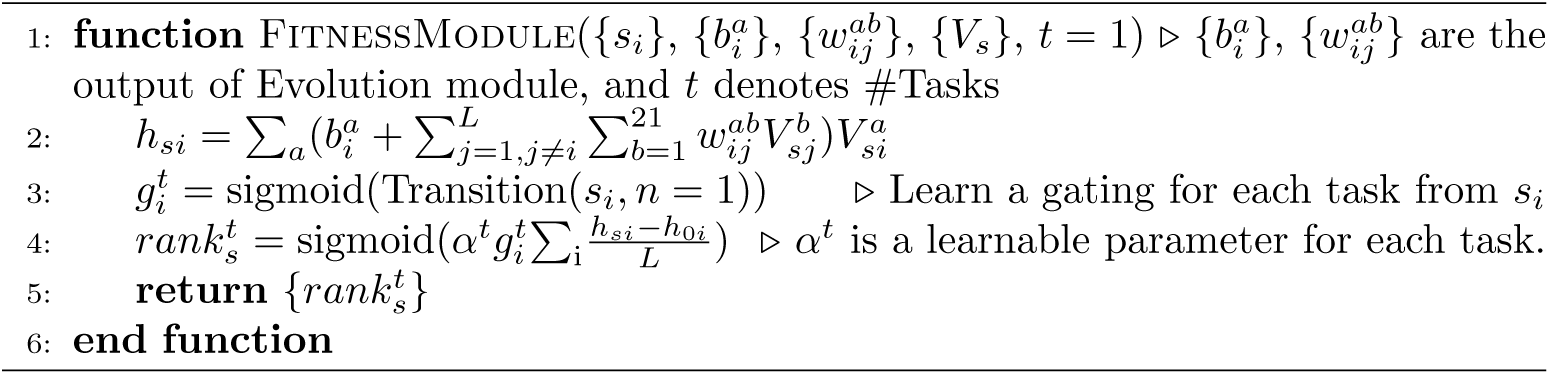

## 2 Supplementary Figures

### 2.1 Receiver-operating characteristic (ROC) curves of the predictions of TCR-pMHC bindings

**Fig. B1.**
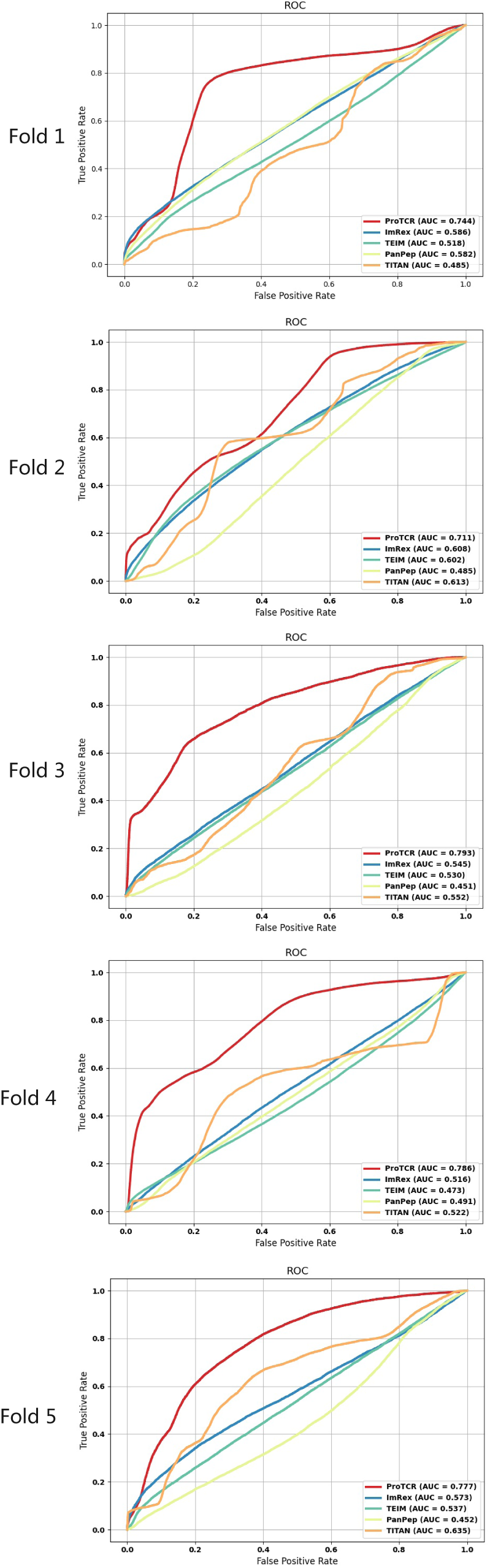
Receiver-operating characteristic (ROC) curves of the predictions by ProTCR, ImRex, TEIM, PanPep, and TITAN. Here, we show the five-fold cross-validation performance of these prediction approaches by drawing the ROC curves for each fold separately

### 2.2 The distribution of peptide-specific AUROC calculated by ProTCR

**Fig. B2.**
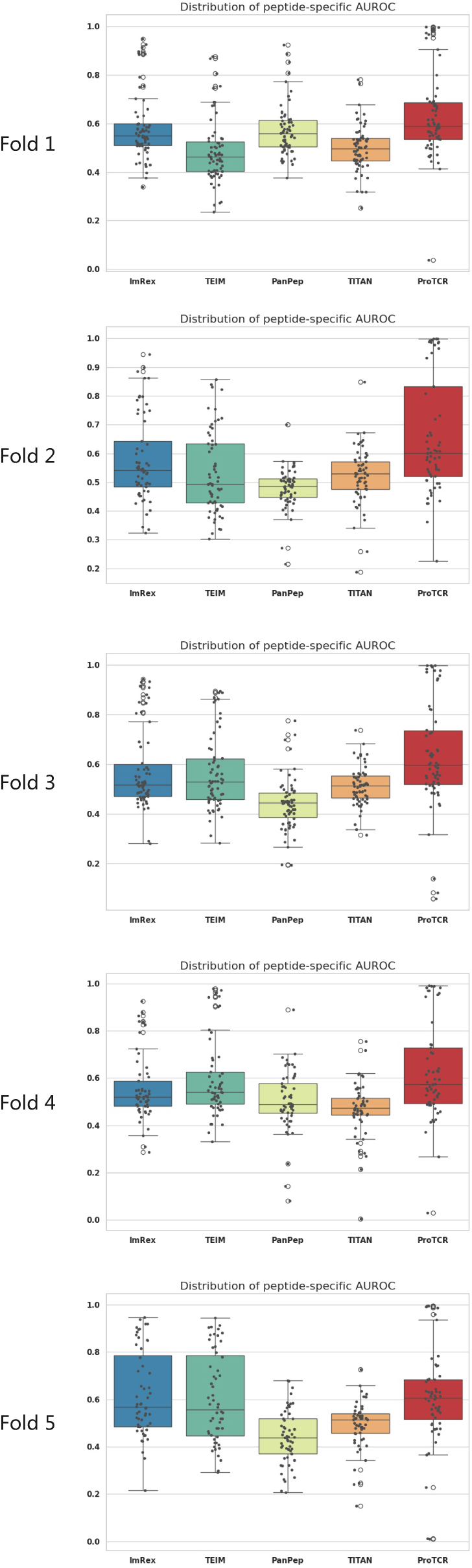
The distribution of peptide-specific AUROC values calculated by ProTCR. obtained from five-fold cross-validation. Here, we show the five-fold cross-validation performance by drawing the distribution of AUROC values for each fold separately

### 2.3 Benchmarking ProTCR against AlphaFold3 on single-cell sequencing test data

**Fig. B3.**
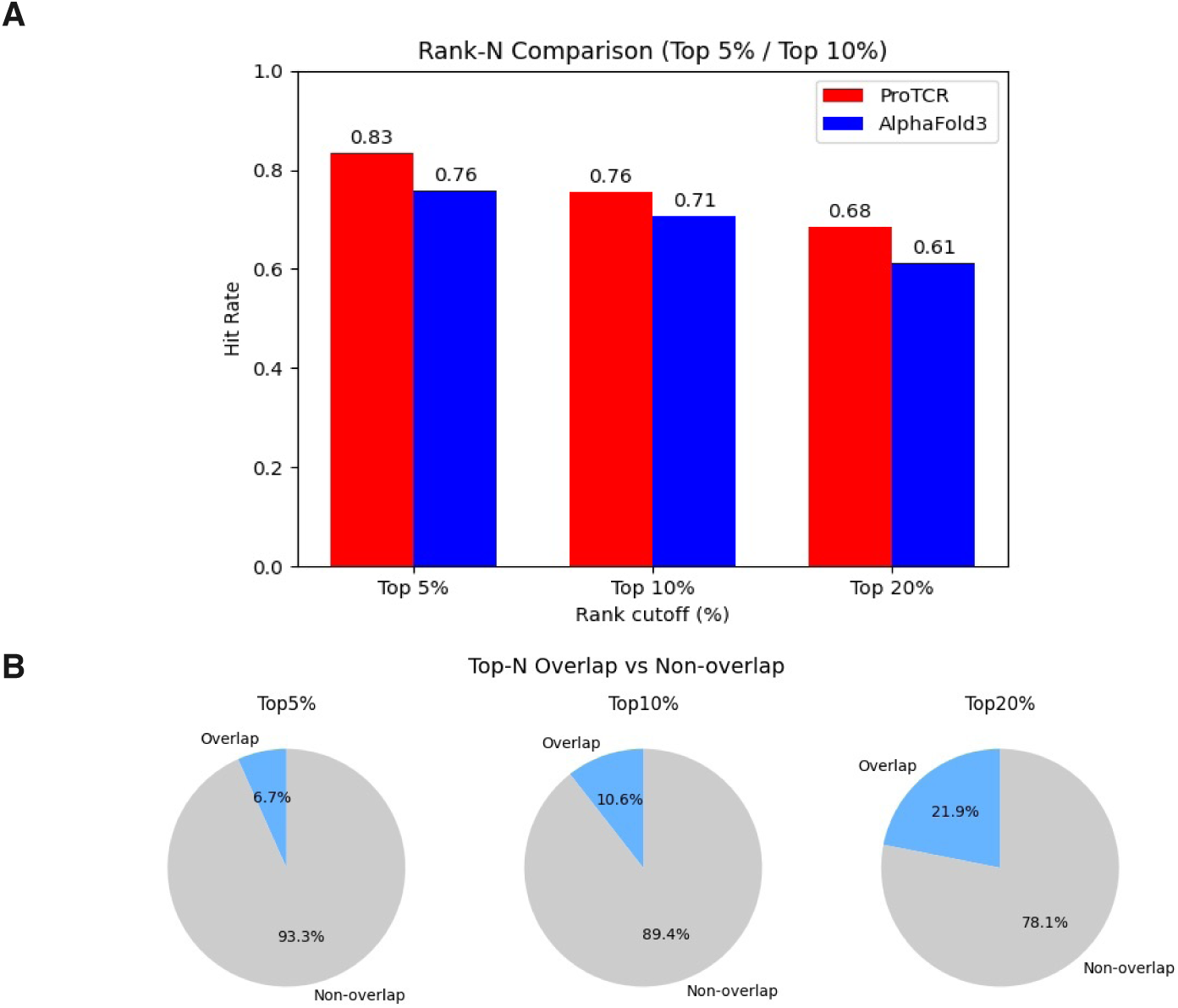
Benchmarking ProTCR against AlphaFold3 on single-cell sequencing test data. A total of 2,000 TCR–pMHC complexes were randomly selected as test set from the 10X Genomics Datasets (https://www.10xgenomics.com/resources/datasets), including 1,000 binding examples and 1,000 non-binding examples.. To prevent data leakage, ProTCR predictions were generated using models trained without the corresponding test entries. A, Positive rates at the top 5%, 10% and 20% ranked predictions, showing that ProTCR outperforms AlphaFold3. B, Overlap of top-ranked predictions between ProTCR and AlphaFold3 is minimal, indicating that the two methods provide complementary information.

### 2.4 Sequence motifs of MHC-specific binding peptides

**Fig. B4.**
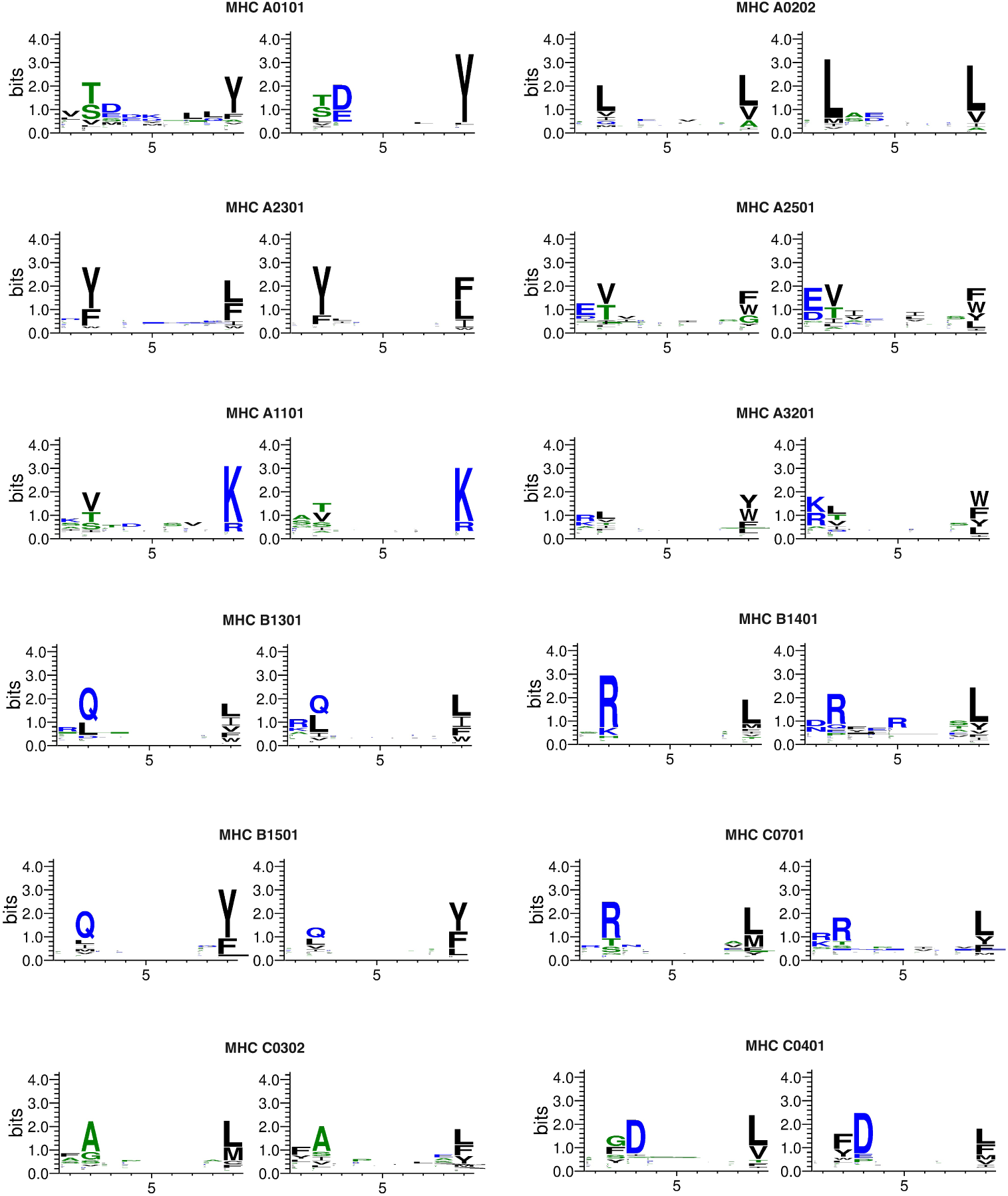
Sequence motifs of MHC-specific binding peptides and the peptides generated by ProTCR for a given MHC. We show the MHC-specific peptides for a total of 12 MHCs, including A0101, A0202, A1101, A2301, A2501, A3201, B1301, B1401, B1501, C0302, C0401, and C0701. These MHC-specific binding peptides exhibit substantial residue preference. The peptides recovered by ProTCR based on MHC and TCR exhibit highly consistent motifs with nearly identical conserved residues.

## 3 Supplementary Data

### 3.1 Supplementary Data 1

Peptide-specific AUROC values for the prediction by ProTCR, ImRex, TEIM, PanPep, and TITAN. We visualize these values in Figure 2B. These values are available from https://github.com/bigict/tcr_pmhc/blob/main/Supplementary/Supplementary_Data1.csv.

### 3.2 Supplementary Data 2

The relationship between AUROC scores and the similarity between COVID-19 pep-tides and those used for training. The Pearson correlation coefficient between AUROC values and peptide similarity is only 0.11, implying that ProTCR can reliably predict TCR-peptide binding even if the peptides are highly distinct from the ones in the train-ing set. We visualize these values in Fig. 5C. These values are available from https://github.com/bigict/tcr_pmhc/blob/main/Supplementary/Supplementary_Data2.csv.

## References

[1] Alex D Waldman, Jill M Fritz, and Michael J Lenardo. A guide to cancer immunotherapy: from T cell basic science to clinical practice. Nature Reviews Immunology, 20(11):651–668, 2020.

[2] Apostolia-Maria Tsimberidou, Karlyle Van Morris, Henry Hiep Vo, Stephen Eck, Yu-Feng Lin, Jorge Mauricio Rivas, and Borje S Andersson. T-cell receptor-based therapy: an innovative therapeutic approach for solid tumors. Journal of Hematology & Oncology, 14:1–22, 2021.

[3] Jamie Rossjohn, Stephanie Gras, John J Miles, Stephen J Turner, Dale I Godfrey, and James McCluskey. T cell antigen receptor recognition of antigen-presenting molecules. Annual Review of Immunology, 33(1):169–200, 2015.

[4] F Stephen Hodi, Steven J O’Day, David F McDermott, Robert W Weber, Jeffrey A Sosman, John B Haanen, Rene Gonzalez, Caroline Robert, Dirk Schaden-dorf, Jessica C Hassel, et al. Improved survival with ipilimumab in patients with metastatic melanoma. New England Journal of Medicine, 363(8):711–723, 2010.

[5] Jedd D Wolchok, Vanna Chiarion-Sileni, Rene Gonzalez, Piotr Rutkowski, Jean-Jacques Grob, C Lance Cowey, Christopher D Lao, John Wagstaff, Dirk Schadendorf, Pier F Ferrucci, et al. Overall survival with combined nivolumab and ipilimumab in advanced melanoma. New England Journal of Medicine, 377(14):1345–1356, 2017.

[6] Weijun Zhou, Jinyi Yu, Yilu Li, and Kankan Wang. Neoantigen-specific TCR-T cell-based immunotherapy for acute myeloid leukemia. Experimental Hematology & Oncology, 11(1):100, 2022.

[7] Aaron P Rapoport, Edward A Stadtmauer, Gwendolyn K Binder-Scholl, Olga Goloubeva, Dan T Vogl, Simon F Lacey, Ashraf Z Badros, Alfred Garfall, Bren-dan Weiss, Jeffrey Finklestein, et al. NY-ESO-1–specific TCR–engineered T cells mediate sustained antigen-specific antitumor effects in myeloma. Nature Medicine, 21(8):914–921, 2015.

[8] Pradyot Dash, Andrew J Fiore-Gartland, Tomer Hertz, George C Wang, Shalini Sharma, Aisha Souquette, Jeremy Chase Crawford, E Bridie Clemens, Thi HO Nguyen, Katherine Kedzierska, et al. Quantifiable predictive features define epitope-specific T cell receptor repertoires. Nature, 547(7661):89–93, 2017.

[9] John-William Sidhom, H Benjamin Larman, Drew M Pardoll, and Alexander S Baras. DeepTCR is a deep learning framework for revealing sequence concepts within T-cell repertoires. Nature Communications, 12(1):1605, 2021.

[10] Hongyi Zhang, Xiaowei Zhan, and Bo Li. GIANA allows computationally-efficient TCR clustering and multi-disease repertoire classification by isometric transformation. Nature Communications, 12(1):4699, 2021.

[11] Hongyi Zhang, Longchao Liu, Jian Zhang, Jiahui Chen, Jianfeng Ye, Sachet Shukla, Jian Qiao, Xiaowei Zhan, Hao Chen, Catherine J Wu, et al. Investiga-tion of antigen-specific T-cell receptor clusters in human cancers. Clinical Cancer Research, 26(6):1359–1371, 2020.

[12] Jacob Glanville, Huang Huang, Allison Nau, Olivia Hatton, Lisa E Wagar, Florian Rubelt, Xuhuai Ji, Arnold Han, Sheri M Krams, Christina Pettus, et al. Identify-ing specificity groups in the T cell receptor repertoire. Nature, 547(7661):94–98, 2017.

[13] Emmi Jokinen, Jani Huuhtanen, Satu Mustjoki, Markus Heinonen, and Harri Läahdesmäki. Predicting recognition between T cell receptors and epitopes with TCRGP. PLoS Computational Biology, 17(3):e1008814, 2021.

[14] Sofie Gielis, Pieter Moris, Wout Bittremieux, Nicolas De Neuter, Benson Ogunjimi, Kris Laukens, and Pieter Meysman. Detection of enriched T cell epitope specificity in full T cell receptor sequence repertoires. Frontiers in Immunology, 10:2820, 2019.

[15] Giancarlo Croce, Sara Bobisse, Dana Léa Moreno, Julien Schmidt, Philippe Guillame, Alexandre Harari, and David Gfeller. Deep learning predictions of tcr-epitope interactions reveal epitope-specific chains in dual alpha T cells. Nature Communications, 15(1):3211, 2024.

[16] Alessandro Montemurro, Viktoria Schuster, Helle Rus Povlsen, Amalie Kai Bentzen, Vanessa Jurtz, William D Chronister, Austin Crinklaw, Sine R Hadrup, Ole Winther, Bjoern Peters, et al. NetTCR-2.0 enables accurate prediction of TCR-peptide binding by using paired TCRα and β sequence data. Communications Biology, 4(1):1060, 2021.

[17] Yu Zhao, Bing He, Fan Xu, Chen Li, Zhimeng Xu, Xiaona Su, Haohuai He, Yueshan Huang, Jamie Rossjohn, Jiangning Song, et al. DeepAIR: A deep learning framework for effective integration of sequence and 3D structure to enable adaptive immune receptor analysis. Science Advances, 9(32):eabo5128, 2023.

[18] Pieter Moris, Joey De Pauw, Anna Postovskaya, Sofie Gielis, Nicolas De Neuter, Wout Bittremieux, Benson Ogunjimi, Kris Laukens, and Pieter Meysman. Current challenges for unseen-epitope TCR interaction prediction and a new perspective derived from image classification. Briefings in Bioinformatics, 22(4):bbaa318, 2021.

[19] Yicheng Gao, Yuli Gao, Yuxiao Fan, Chengyu Zhu, Zhiting Wei, Chi Zhou, Guo-hui Chuai, Qinchang Chen, He Zhang, and Qi Liu. Pan-peptide meta learning for T-cell receptor–antigen binding recognition. Nature Machine Intelligence, 5(3):236–249, 2023.

[20] Xingang Peng, Yipin Lei, Peiyuan Feng, Lemei Jia, Jianzhu Ma, Dan Zhao, and Jianyang Zeng. Characterizing the interaction conformation between T-cell receptors and epitopes with deep learning. Nature Machine Intelligence, 5(4):395–407, 2023.

[21] Anna Weber, Jannis Born, and María Rodriguez Martínez. TITAN: T-cell receptor specificity prediction with bimodal attention networks. Bioinformatics, 37(Supplement 1):i237–i244, 2021.

[22] Tianshi Lu, Ze Zhang, James Zhu, Yunguan Wang, Peixin Jiang, Xue Xiao, Chantale Bernatchez, John V Heymach, Don L Gibbons, Jun Wang, et al. Deep learning-based prediction of the T cell receptor–antigen binding specificity. Nature Machine Intelligence, 3(10):864–875, 2021.

[23] Jolan Wauters. ERGO-II: An improved Bayesian optimization technique for robust design with multiple objectives, failed evaluations, and stochastic parameters. Journal of Mechanical Design, 146(10):101704, 2024.

[24] Josh Abramson, Jonas Adler, Jack Dunger, Richard Evans, Tim Green, Alexander Pritzel, Olaf Ronneberger, Lindsay Willmore, Andrew J Ballard, Joshua Bambrick, et al. Accurate structure prediction of biomolecular interactions with AlphaFold 3. Nature, 630(8016):493–500, 2024.

[25] John Jumper, Richard Evans, Alexander Pritzel, Tim Green, Michael Figurnov, Olaf Ronneberger, Kathryn Tunyasuvunakool, Russ Bates, Augustin Žídek, Anna Potapenko, et al. Highly accurate protein structure prediction with AlphaFold. nature, 596(7873):583–589, 2021.

[26] Rohith Krishna, Jue Wang, Woody Ahern, Pascal Sturmfels, Preetham Venkatesh, Indrek Kalvet, Gyu Rie Lee, Felix S Morey-Burrows, Ivan Anishchenko, Ian R Humphreys, et al. Generalized biomolecular modeling and design with RoseTTAFold All-Atom. Science, 384(6693):eadl2528, 2024.

[27] Marius Messemaker, Bjørn P.Y. Kwee, Živa Moravec, Daniel Álvarez-Salmoral, Jos Urbanus, Sam de Paauw, Jeroen Geerligs, Rhianne Voogd, Ben Morris, Aurélie Guislain, Maike Mußmann, Yaël Winkler, Maxime Steinmetz, Matyas Iras, Eric Marcus, Jonas Teuwen, Anastassis Perrakis, Roderick L. Beijersbergen, Wouter Scheper, and Ton N. Schumacher. A functionally validated TCR-pMHC database for TCR specificity model development. bioRxiv, 2025.

[28] LOUIS A Matis, ANGEL Ezquerra, and JOHN E Coligan. Expression of two distinct T cell receptor alpha/beta heterodimers by an antigen-specific T cell clone. The Journal of Experimental Medicine, 168(6):2379–2384, 1988.

[29] David S Fischer, Yihan Wu, Benjamin Schubert, and Fabian J Theis. Predicting antigen specificity of single T cells based on TCR CDR3 regions. Molecular Systems Biology, 16(8):e9416, 2020.

[30] Garry Dolton, Katie Tungatt, Angharad Lloyd, Valentina Bianchi, Sarah M Theaker, Andrew Trimby, Christopher J Holland, Marco Donia, Andrew J God-kin, David K Cole, et al. More tricks with tetramers: a practical guide to staining T cells with peptide–MHC multimers. Immunology, 146(1):11–22, 2015.

[31] John D Altman, Paul AH Moss, Philip JR Goulder, Dan H Barouch, Michael G McHeyzer-Williams, John I Bell, Andrew J McMichael, and Mark M Davis. Phe-notypic analysis of antigen-specific T lymphocytes. Science, 274(5284):94–96, 1996.

[32] Ido Springer, Nili Tickotsky, and Yoram Louzoun. Contribution of T cell receptor alpha and beta CDR3, MHC typing, V and J genes to peptide binding prediction. Frontiers in Immunology, 12:664514, 2021.

[33] Pieter Moris, Joey De Pauw, Anna Postovskaya, Sofie Gielis, Nicolas De Neuter, Wout Bittremieux, Benson Ogunjimi, Kris Laukens, and Pieter Meysman. Cur-rent challenges for unseen-epitope TCR interaction prediction and a new perspec-tive derived from image classification. Briefings in Bioinformatics, 22(4):bbaa318, 2021.

[34] Filippo Grazioli, Anja Mösch, Pierre Machart, Kai Li, Israa Alqassem, Timothy J O’Donnell, and Martin Renqiang Min. On TCR binding predictors failing to generalize to unseen peptides. Frontiers in Immunology, 13:1014256, 2022.

[35] Caitlin D Castro, Adrienne M Luoma, and Erin J Adams. Coevolution of T-cell receptors with MHC and non-MHC ligands. Immunological Reviews, 267(1):30–55, 2015.

[36] Sneha Rangarajan and Roy A Mariuzza. T cell receptor bias for MHC: co-evolution or co-receptors? Cellular and Molecular Life Sciences, 71(16):3059–3068, 2014.

[37] Josh Achiam, Steven Adler, Sandhini Agarwal, Lama Ahmad, Ilge Akkaya, Florencia Leoni Aleman, Diogo Almeida, Janko Altenschmidt, Sam Altman, Shyamal Anadkat, et al. GPT-4 technical report. arXiv preprint arXiv:2303.08774, 2023.

[38] Kevin E Wu, Kathryn Yost, Bence Daniel, Julia Belk, Yu Xia, Takeshi Egawa, Ansuman Satpathy, Howard Chang, and James Zou. TCR-BERT: learning the grammar of T-cell receptors for flexible antigen-binding analyses. In Machine Learning in Computational Biology, pages 194–229. PMLR, 2024.

[39] Kamilla Kjærgaard Jensen, Vasileios Rantos, Emma Christine Jappe, Tobias Hegelund Olsen, Martin Closter Jespersen, Vanessa Jurtz, Leon Eyrich Jessen, Esteban Lanzarotti, Swapnil Mahajan, Bjoern Peters, et al. TCRpMHC-models: Structural modelling of TCR-pMHC class I complexes. Scientific Reports, 9(1):14530, 2019.

[40] Xinyi Xu, Hua Li, and Chenqi Xu. Structural understanding of T cell receptor triggering. Cellular & Molecular Immunology, 17(3):193–202, 2020.

[41] Leslie Pack Kaelbling, Michael L Littman, and Andrew W Moore. Reinforcement learning: A survey. Journal of Artificial Intelligence Research, 4:237–285, 1996.

[42] Ameer Haj-Ali, Nesreen K Ahmed, Ted Willke, Yakun Sophia Shao, Krste Asanovic, and Ion Stoica. Neurovectorizer: End-to-end vectorization with deep reinforcement learning. In Proceedings of the 18th ACM/IEEE International Symposium on Code Generation and Optimization, pages 242–255, 2020.

[43] Rafael Rafailov, Archit Sharma, Eric Mitchell, Christopher D Manning, Stefano Ermon, and Chelsea Finn. Direct preference optimization: Your language model is secretly a reward model. Advances in Neural Information Processing Systems, 36:53728–53741, 2023.

[44] Randi Vita, Swapnil Mahajan, James A Overton, Sandeep Kumar Dhanda, Sheridan Martini, Jason R Cantrell, Daniel K Wheeler, Alessandro Sette, and Bjoern Peters. The immune epitope database (IEDB): 2018 update. Nucleic Acids Research, 47(D1):D339–D343, 2019.

[45] Dmitry V Bagaev, Renske MA Vroomans, Jerome Samir, Ulrik Stervbo, Cristina Rius, Garry Dolton, Alexander Greenshields-Watson, Meriem Attaf, Evgeny S Egorov, Ivan V Zvyagin, et al. VDJdb in 2019: database extension, new analysis infrastructure and a T-cell receptor motif compendium. Nucleic Acids Research, 48(D1):D1057–D1062, 2020.

[46] Yuepeng Jiang, Miaozhe Huo, and Shuai Cheng Li. TEINet: a deep learning framework for prediction of TCR-epitope binding specificity. Briefings in Bioinformatics, 24(2):bbad086, 2023.

[47] David F Burke, Patrick Bryant, Inigo Barrio-Hernandez, Danish Memon, Gabriele Pozzati, Aditi Shenoy, Wensi Zhu, Alistair S Dunham, Pascal Albanese, Andrew Keller, et al. Towards a structurally resolved human protein interaction network. Nature Structural & Molecular Biology, 30(2):216–225, 2023.

[48] Birkir Reynisson, Bruno Alvarez, Sinu Paul, Bjoern Peters, and Morten Nielsen. NetMHCpan-4.1 and NetMHCIIpan-4.0: improved predictions of MHC antigen presentation by concurrent motif deconvolution and integration of MS MHC eluted ligand data. Nucleic Acids Research, 48(W1):W449–W454, 2020.

[49] John-William Sidhom and Alexander S Baras. Analysis of SARS-CoV-2 specific T-cell receptors in ImmuneCode reveals cross-reactivity to immunodominant Influenza M1 epitope. BioRxiv, pages 2020–06, 2020.

[50] Huang Huang, Michael J Sikora, Saiful Islam, Roshni Roy Chowdhury, Yueh-hsiu Chien, Thomas J Scriba, Mark M Davis, and Lars M Steinmetz. Select sequencing of clonally expanded CD8+ T cells reveals limits to clonal expansion. Proceedings of the National Academy of Sciences, 116(18):8995–9001, 2019.

[51] Ming Li, Ngoc Hieu Tran, Chao Peng, Qingyang Lei, Lei Xin, Jingxiang Lang, Qing Zhang, Wenting Li, Rui Qiao, Haiming Qin, et al. A complete mass spectrometry-based immunopeptidomics pipeline for neoantigen identification and validation. form https://assets-eu.researchsquare.com/files/rs-3520814/v1coveredc6f84eec-8166-4e7c-b8a3-cadf4c594bb5.pdf?c=1702920854, 2023.

[52] L Steven Johnson, Sean R Eddy, and Elon Portugaly. Hidden Markov model speed heuristic and iterative HMM search procedure. BMC Bioinformatics, 11:1–8, 2010.

[53] Trupti M Kodinariya, Prashant R Makwana, et al. Review on determining number of Cluster in K-Means Clustering. International Journal, 1(6):90–95, 2013.

